# The complete chloroplast genome of Chinese medicine (*Psoralea corylifolia Linn*): Molecular Structures, barcoding analysis, and phylogenetic Analysis

**DOI:** 10.1101/587758

**Authors:** Wei Tan, Han Gao, Huanyu Zhang, Xiaolei Yu, Xiaoxuan Tian, Weiling Jiang, Kun Zhou

## Abstract

*Psoralea corylifolia* is one kind of traditional Chinese medicine used in China widely. In this study, we sequence the complete chloroplast genome of *P. corylifolia*, which is 153,114 bp in size and includes a pair of inverted repeats regions of 25,557 bp interspersed by a small single copy of 17,885 bp and a large single copy of 84,115 bp region. Approximately 98 simple sequence repeats, 14 forward, 2 reverse, 2 complement, 32 palindromic and 49 tandem repeats are identified in the *P. corylifolia* chloroplast genome. The chloroplast genomes of *P. corylifolia* and three *Glycine* species are conserved in gene order and content, but show high diversity within intergenic spacers. *P. corylifolia* with three *Glycine* species in Papilionoideae fall into the same clade based on 75 conserved coding-protein genes phylogenomic analysis. Moreover, four chloroplast DNA regions (*ycf1, matK, accD, ndhF*) can serve as the barcodes. In general, our findings will dedicate to better comprehension of the genome aspect as well as evolutionary status of *P. corylifolia.*

## Introduction

Leguminosae is known as the third-largest family of angiosperm in terms of 751 genera, consisting of 19,000 species[1, 2]. The subfamily Papilionoideae of Leguminosae compose of several tribes, including Fabeae, Galegeae, Indigofereae, Loteae, Millettieae, Phaseoleae, Psoraleeae, *etc*[3]. Among these tribes, nucleotide substitution rates of Psoraleeae have been found elevated relative to other tribes, suggestive of rapid evolution or diversification[4].The genus *Psoralea of* Psoraleeae contains about 120 species mainly distributed in southern Africa, North and South America, Australia and only one species lives in China[5]. *Psoralea Corylifolia* (Chinese name Buguzhi) is an annual plant, generating pale-purple flowers. Since the occurrence of minute brown glands, the plant has a distinctive and pleasant fragrance[6]. Furthermore, the seed of *P. Corylifolia*, which can traditionally be used for the treatment of menopause, depression, kidney deficiency[7, 8]. Previous pharmacological studies indicate its antioxidant, antimicrobial, antiinflammatory and chemoprotective properties. Thus, *P. Corylifolia* has been widely used in traditional Chinese medicine for its pharmacological effects to treat multiple diseases[9]. Based on its medicinal value, *P. Corylifolia* deserves utilization and development. Recently, with the rapid growth of next-generation sequencing (NGS) technologies, more and more genomic resources at reasonable schedules and prices have been provided[10]. However, despite *P. Corylifolia* is very important, the studies regarding its genetic character are scarce. Therefore, the genetic variety and phylogenetic status of this species needs analyzing by molecular techniques.

Chloroplast genomes, which comprise a typical structure consisting of two duplicate inverted repeat (IRs) isolated by the large and small single copy (LSC and SSC) regions, frequently have conserved annular double-stranded structure, ranging from 120 to 160 kb in length[11, 12]. However, previous studies have revealed that the chloroplast genomes have shown notable structural variation among Papilionoideae subfamily[13]. Meanwhile, most species in Papilionoideae possess a large (50-kb) inversion in their chloroplast genomes[14]. Not only that, the loss of IRs have been detected in many species of Papilionoideae, such as *Glycyrrhiza glabra, Lens culinaris, Vicia faba*[15]. The genes obtained from the chloroplast genome have been applied to molecular identification and phylogenetic evolution analysis, as a result of its maternal inheritance pattern and comparatively independent evolution[16]. Besides that, although previous studies have recommended some loci as the plant barcode, including *rbcL, matK, trnH-psbA, trnL-trnF*[17, 18], the complete chloroplast genomes might be properer to be used as a super-barcode for species identification[19].

For such an important group, the classification and phylogenetic relationships of Psoraleeae remain poorly known, even no chloroplast genomes of this tribe have been reported. Therefore, in this research, we confirm the first complete chloroplast genome sequence of *Psoralea Corylifolia* and compare *P. Corylifolia* with the published chloroplast genome of the related genus of *Glycine*, including *Glycine max*[20], *Glycine soja*[21] and *Glycine gracilis*[22]. Our aim is to construct and characterize the structure of the complete chloroplast genome of *Psoralea Corylifolia* and provide vital phylogenetic and genetic information for future studies in Psoraleeae and legume plastomes.

## Materials and methods

### Chloroplast genome sequencing and assembly

The standard sample was bought from National Institutes for Food and Drug Control, which batch number was 121056-200904. And the voucher was deposited in Tianjin State Key Laboratory of Modern Chinese Medicine. Total genomic DNA was detached using Extract Genomic DNA Kit following the protocol of the manufacturer. Then, Illumina Animal and Plant Small Fragments Libraries were prepared from the sample with an average insert size of 800 bp. The libraries were sequenced with 2 × 150 bp on the Hiseq instrument. The quality of raw data was assessed with FastQC. The complete chloroplast genome of *Glycine soja* (GenBank Accession: NC_022868) was used as seed in the assembly of *Psoralea corylifolia* and the paired-end reads were assembled by NOVOPlasty 2.6.3[23] with k-mer length 39. The contigs were subsequently oriented to construct the complete chloroplast genome by mapping to reference (NC_022868). Where required, the raw data was mapped to the chloroplast genome with CLC workbench v11.0.1 to resolve degenerate bases. We used DOGMA[24] (http://dogma.ccbb.utexas.edu/) to annotate the complete plastid genome, and besides that, tRNAscan-SE[25] (http://lowelab.ucsc.edu/tRNAscan-SE/) was used to confirm the annotations of tRNA. Finally, all of the annotations were carefully manually checked against the reference (NC_022868, NC_030329, NC_007942), and confirmed in Sequin v15.50. OGDRAW[26] (https://chlorobox.mpimp-golm.mpg.de/OGDraw.html) was used to create the physical map of the circular chloroplast genome of this species. The annotated chloroplast genome sequence of *Psoralea corylifolia* was deposited in GenBank (Accession Number: MK069582).

### Comparative genome analysis

The complete chloroplast genome of *Psoralea corylifolia* was compared with three species (*Glycine max, Glycine gracilis, Glycine soja*), using the mVISTA[27] (http://genome.lbl.gov/vista/mvista/submit.shtml) program in a LAGAN mode, while *Glycine max* was set as a reference. IRscope[28] (https://irscope.shinyapps.io/irapp/) was used to visualize the genes on the boundaries of LSC, SSC, and IRs according to their annotations.

### Codon usage

Codon usage was determined for all protein-coding genes, the relative synonymous codon usage (RSCU) was calculated using MEGA X[29]. Besides, the ggplot2 package was used to visualize codon usage frequency.

### Analysis of single sequence repeats and tandem repeats

REPuter[30] (http://bibiserv2.cebitec.unibielefeld.de/reputer?id=reputer_view_submission) was used to identify forward, reverse, complement and palindromic repeats in *Psoralea corylifolia* chloroplast genome. The minimal repeat size was set as 30 bp and the Hamming distance was 3. In addition to REPuter, MISA[31] was used to detect single sequence repeats (SSR), with the following setting: >10 for mononucleotide, >5 for dinucleotide, >4 for trinucleotide, and >3 for tetra-nucleotide, pentanucleotide, and hexanucleotide SSRs. Furthermore, Tandem Repeats Finder[32] (http://tandem.bu.edu/trf/trf.submit.options.html) was used to find tandem repeats with the default setting.

### RNA editing site prediction

The online program Predictive RNA Editor for Plants (PREP) suite[33] (http://prep.unl.edu/) was used to predict potential RNA editing sites using 35 protein-coding genes (*matK, atpB, ycf3, psaB, rps14, rpoB, rpoC1, rpoC2, rps2, atpI, atpF, atpA, rps16, accD, psaI, psbL, psbF, psbE, petL, petG, rpl20, clpP, psbB, petB, petD, rpoA, rps8, rpl2, rpl23, ndhB, ndhA, ndhG, ndhD, ccsA, ndhF*) of *Psoralea corylifolia* chloroplast genome with default parameters.

### Synonymous and nonsynonymous substitution rate analysis

The genes of the species (*P. corylifolia, G. max, G. gracilis,* and *G. soja*) with the same functions were grouped following previous studies[34-36]. The singular genes (*clpP, cemA, matK, ccsA*, and *ycf1)* and the same functions genes such as ATP synthase (*atp*), Cytochrome b6/f complex (*pet*), Subunits of NADH-dehydrogenase (*ndh*), Photosystem I (*psa*), Photosystem II (*psb*), Large subunit of ribosomal (*rpl*), DNA-dependent RNA polymerase (*rpo*) and Small subunit of ribosomal proteins (*rps*) were analyzed. PAML v4.9 package[37] was used to analyze synonymous and nonsynonymous substitution rate, and *Psoralea corylifolia* was set as outgroup. Within the PAML package, the yn00 program was utilized to calculate dN, dS, and dN/dS. Each sequence was set as 3×4 (codon position×base) table, and we used the Nei–Gojobori (1986) method[38] to confirm the value of dN/dS.

### Divergent Hotspots

MAFFT v7[39] was applied to align the four complete chloroplast genomes. Furthermore, the nucleotide polymorphism of the chloroplast genomes were estimated by DnaSP v6.11.01[40]. The following conditions for sliding window option were set as: window length of 600 bp, step size of 200 bp.

### Phylogenetic and DNA barcoding analysis

Previous studies indicated that the chloroplast genome was qualified to solve the phylogenetic relationship of Fabaceae[13]. In this study, we compared 36 species of Papilionoideae, including *P. corylifolia, G.max, G.gracilis, G.soja*. With the exception of these four species, the remaining 32 species were downloaded from the GenBank (S1 Table). It’s worth noting that the 36 species represented 36 genera in Papilionoideae subfamily. The 75 protein-coding genes were extracted manually and aligned separately using MAFFT v7, and then these matrices were concatenated to form a final data matrix. The phylogenetic relationship of Papilionoideaea was rebuilt while *Pararchidendron pruinosum* was set as the outgroup. RAxML version 8.2.4[41] was used to perform Maximum likelihood (ML) analysis, using 1000 replicates of rapid bootstrap with the GTRGAMMAI substitution model. Bayesian inference (BI) was implemented using MrBayes v3.2.6[42] with default parameters (Lset nst=6, rates=invgamma, Prset statefreqpr=dirichlet(1,1,1,1)). In our study, we found Psoraleeae was more closely related to the Phaseoleae than to the other tribes in the Papilionoideae. Therefore, 9 species of two tribes were collected to test the potential barcodes. Four various regions in the chloroplast genomes have been proposed as DNA barcodes, including *ycf1, matK, accD, ndhF,* genes. The ML trees were reconstructed using the four genes with GTRGAMMAI model, respectively.

## Results and discussion

### Chloroplast genome assembly

By using the Illumina sequencing platform, we obtained a total of 12,330,274 reads with an average read length of 150 bp. The reads were re-mapped to the chloroplast genome, and the coverage of the chloroplast genome was 3,241×. The size of the whole genome was 153,114 bp.

### Organization and gene content

The complete plastid genome of *Psoralea corylifolia* displayed a typical quadripartite structure (Fig 1), which comprised of a pair of inverted repeats (each 25,557 bp in length) separated by the SSC and the LSC region (17,885 and 84,115 bp in length, respectively). All the above values were similar to three *Glycine* species which can be seen in the table (Table 1). The GC content of the complete chloroplast genome was 35.5%. The GC content of the IR (41.9%) was higher than that in LSC (32.7%) and SSC regions (28.8%). The complete chloroplast genome consisted of 111 unique genes, including 77 protein-coding genes, 30 tRNA genes, 4 rRNA genes. Among these genes, 16 genes (*petB, petD, rpl2, rpl16, rps16, rpoC1, trnI-GAU, trnA-UGC, trnG-UCC, trnL-UAA, trnV-UAC, trnK-UUU, ndhA, ndhB, rps12, atpF, clpP* and *ycf3*) included one intron, excepted *ycf3* and *clpP* genes who contained two. In the functional classification of genes, 39 genes were involved in photosynthesis, 5 for Photosystem I, 15 for Photosystem II, 6 for Cytochrome b6/f complex, 6 for different subunits of ATP synthase and 1 for RuBisCO. 5 genes were associated with different functions, and besides, the function of *ycf* genes (*ycf1, ycf2, ycf3*) existing in *Psoralea corylifolia* was unknown (Table 2).

**Table 1.** Characteristics of the chloroplast genomes in four species.

| Species | *P.corylifolia* | *G.max* | *G.gracilis* | *G.soja* |
| --- | --- | --- | --- | --- |
| Size (bp) | 153,114 | 152,218 | 152,218 | 152,217 |
| LSC (bp) | 84,115 | 83,175 | 83,175 | 83,174 |
| IR (bp) | 25,557 | 25,574 | 25,574 | 25,574 |
| SSC (bp) | 17,885 | 17,895 | 17,895 | 17,895 |
| genes | 111 | 111 | 111 | 111 |
| CDS | 77 | 77 | 77 | 77 |
| tRNA | 30 | 30 | 30 | 30 |
| rRNA | 4 | 4 | 4 | 4 |
| GC% | 35.3% | 34% | 35.37% | 35.4% |

**Table 2.**
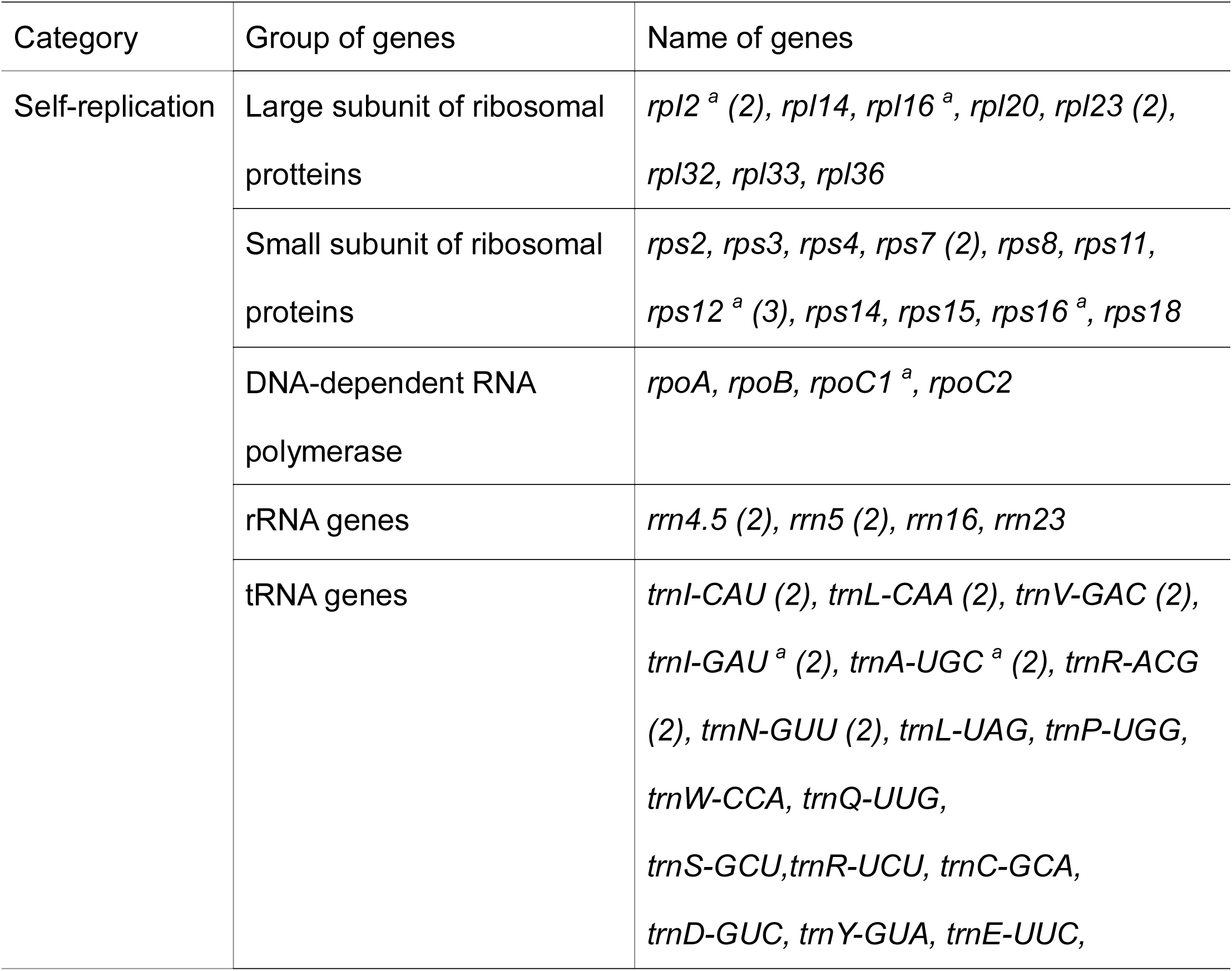
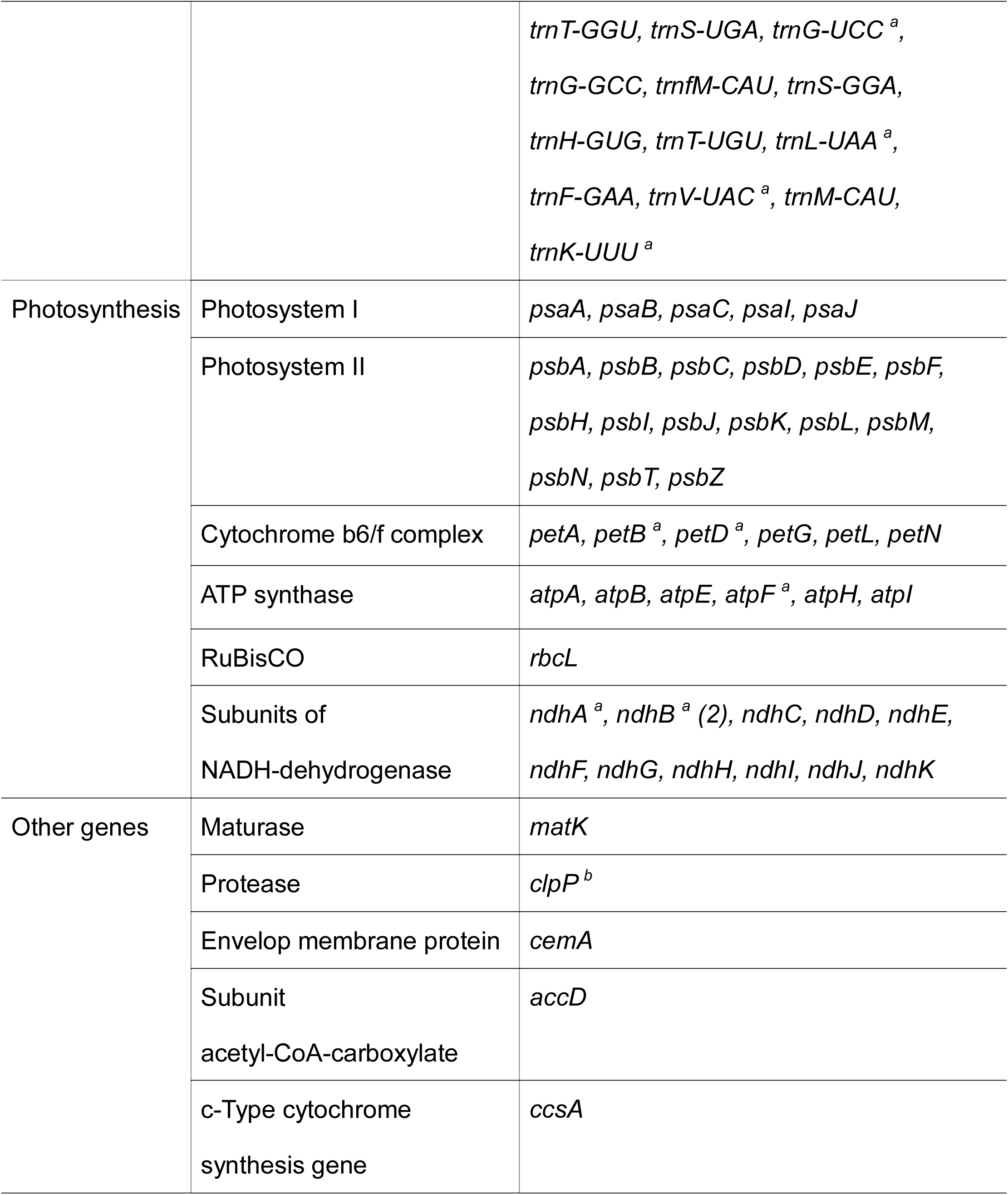
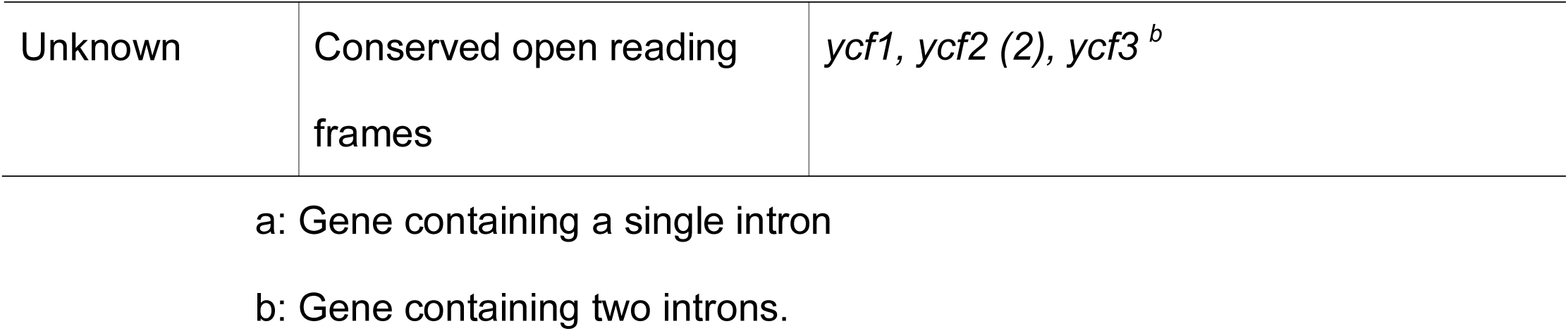
Genes in the sequenced *Psoralea corylifolia* genome.

**Fig 1.**
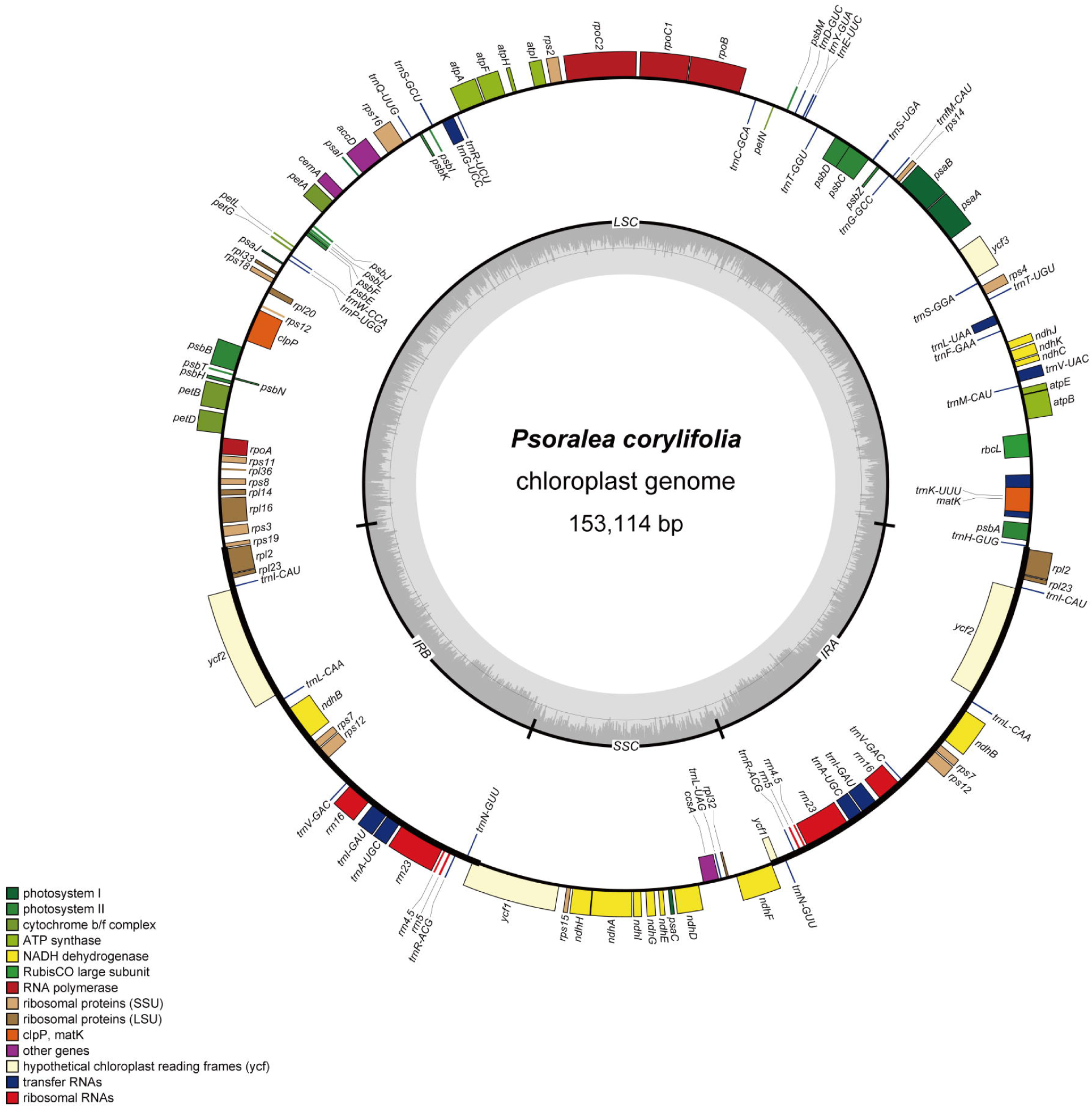
Physical map of the chloroplast genome of the *Psoralea corylifolia.* The genes drawn inside and outside of the circle are transcribed clockwise and counterclockwise, respectively. Different colors represented different functional genes. The GC content of the genome is plotted on the inner circle as dark gray, whereas the light gray corresponded to AT content.

### Comparative genome analysis

The mVISTA program was utilized to analyze the whole sequence identity of the four chloroplast genomes, with *G.max* as a reference. As expected, the alignment of the chloroplast genome showed high sequence similarity, which indicated that they were highly conserved at the scale of genome level. The IR regions were less divergent than LSC and SSC regions, and besides that, more sequence variations were detected to be located in the non-coding regions rather than coding regions. Furthermore, the divergent coding regions were within *ycf1, matK, accD, ndhF*, which could be used as specific DNA barcodes. The *trnH-psbA, trnK-rbcL, trnG-psbZ, psbD-trnT-trnE, petN-trnC, trnG-trnS, trnR-trnG-trnS, ndhI-ndhG, trnL-rpl32* loci and *petD, rpl16, ndhA* introns were most divergent regions, which were located in the intergenic spacers (IGSs) and introns, respectively. The boundary of IR regions indicated hypervariable regions with many nucleotide changes in chloroplast genomes of closely related species[43], simultaneously, the IR border expansion/contraction was considered as an evolutionary event[36]. In this study, we compared genes on the boundaries of the junction sites of the *Psoralea corylifolia* with other three species of *Glycine* (*Glycine max, Glycine gracilis, Glycine soja*) (Fig 2). The IRb/LSC (JLB) border of *Psoralea corylifolia* was located on *rps19* gene, with a size of pseudogene as 61 bp, showing the length 7 bp shorter than other species. The *ycf1* gene crossed the IRb/SSC (JSB) region, and the size of the *ycf1* pseudogene was 5,378 bp in all three *Glycine* species and 5,405 bp in *P. corylifolia*. The SSC/IRA (JSA) border extended into *ndhF* genes in *P. corylifolia* by 9 bp, whereas the *ndhF* genes of the other 3 species were all found in the SSC region. The *trnH-GUG* gene was at the IRa/LSC (JLA) border among the three *Glycine* species and *P. corylifolia* by 2 bp and 16 bp, respectively (Fig 3).

**Fig 2.**
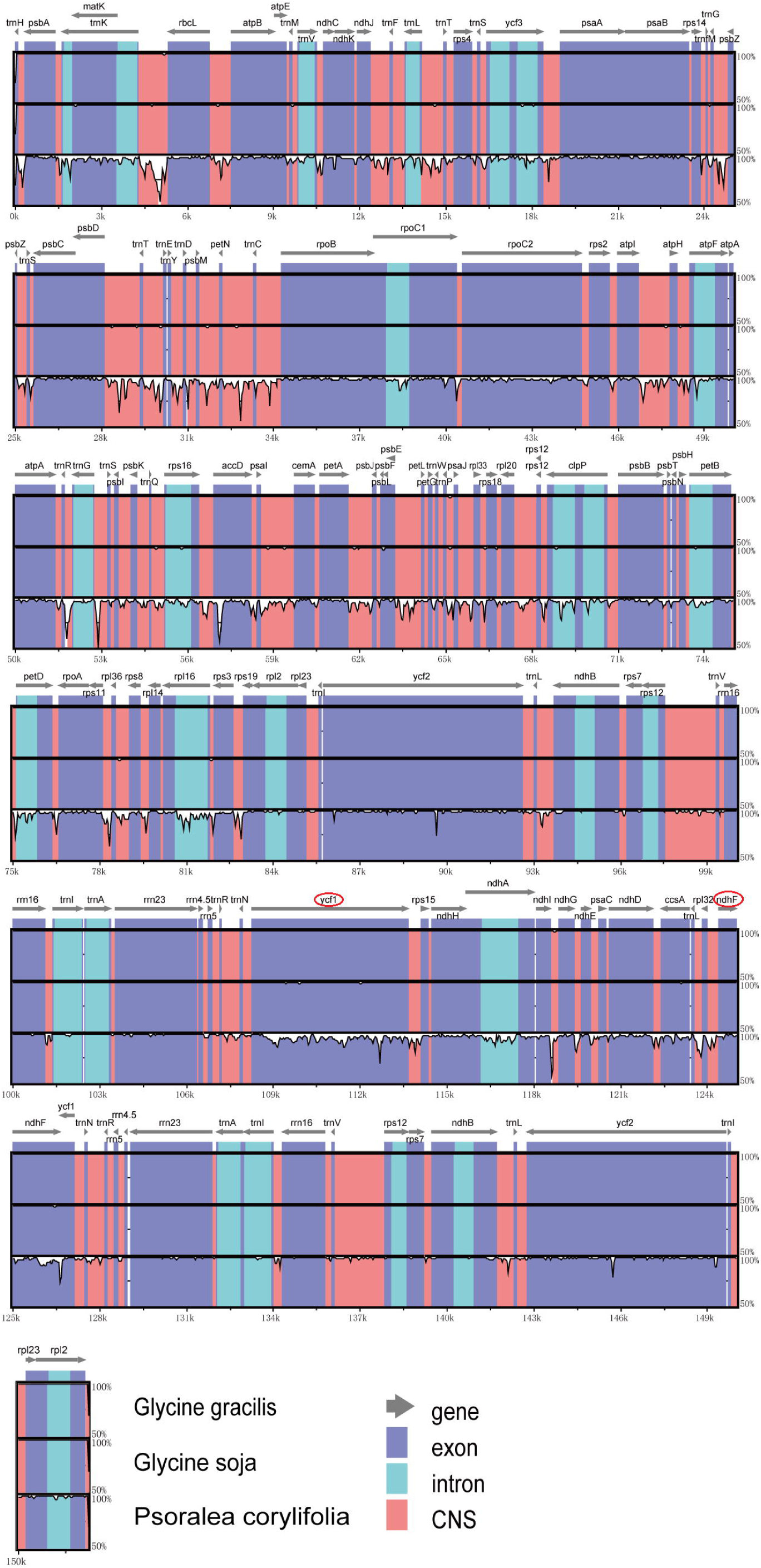
Sequence identity plots among chloroplast genomes of *P. corylifolia, G. max, G. gracilis, G. soja, with G. max* as a reference. Grey arrows above the alignment indicate the position and orientation of genes. The vertical scale represents the percent identity among 50-100%. Exon, intron, and conserved non-coding sequences (CNS) are colored purple, blue and red, respectively.

**Fig 3.**
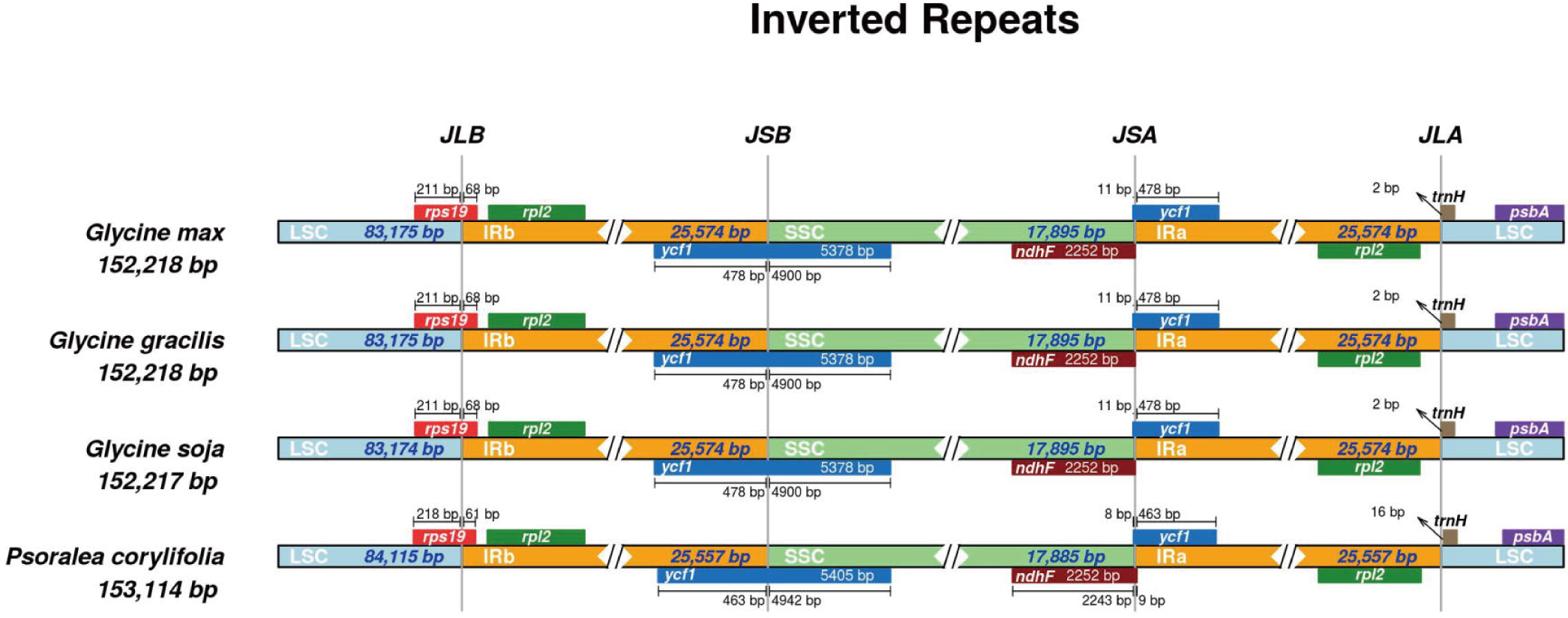
Comparison of LSC, IR, and SSC junction positions among chloroplast genomes of *P. corylifolia, G. max, G. gracilis, G. soja*. The JLB, JSB, JSA and JLA refer to junctions of IRb/LSC, IRb/SSC, SSC/IRa and IRa/LSC, respectively.

### Codon usage frequency analysis

The codon usage frequency was generally linked with gene function both in the level of individual amino acids and nucleotides[44], and besides that, previous evidence indicated that the chloroplast genome might display particular features of codon usage[45]. Based on the nucleic acid sequence in the protein-coding region, MEGA X was used to calculate codon usage frequency and RSCU in the *P. corylifolia* chloroplast genome. The protein-coding genes were composed of 77,916 bp which encoded 25,972 codons. In addition, the most frequently used amino acids were leucine encoded 2,747 (10.6%), whereas only 306 (1.2%) encoded cysteine was the least frequently used amino acids. Almost all the chloroplast genomes strongly biased A-and/or U-ending codons. The result also revealed that all bias favoring synonymous codons (RSCU > 1) with an A or U at the third codon position except for UUG, CUA, UGA. Usage of the initial codon AUG and tryptophan UGG, liked other land plant chloroplast genomes, demonstrated no bias (RSCU = 1) (S2 Table) (Fig 4).

**Fig 4.**
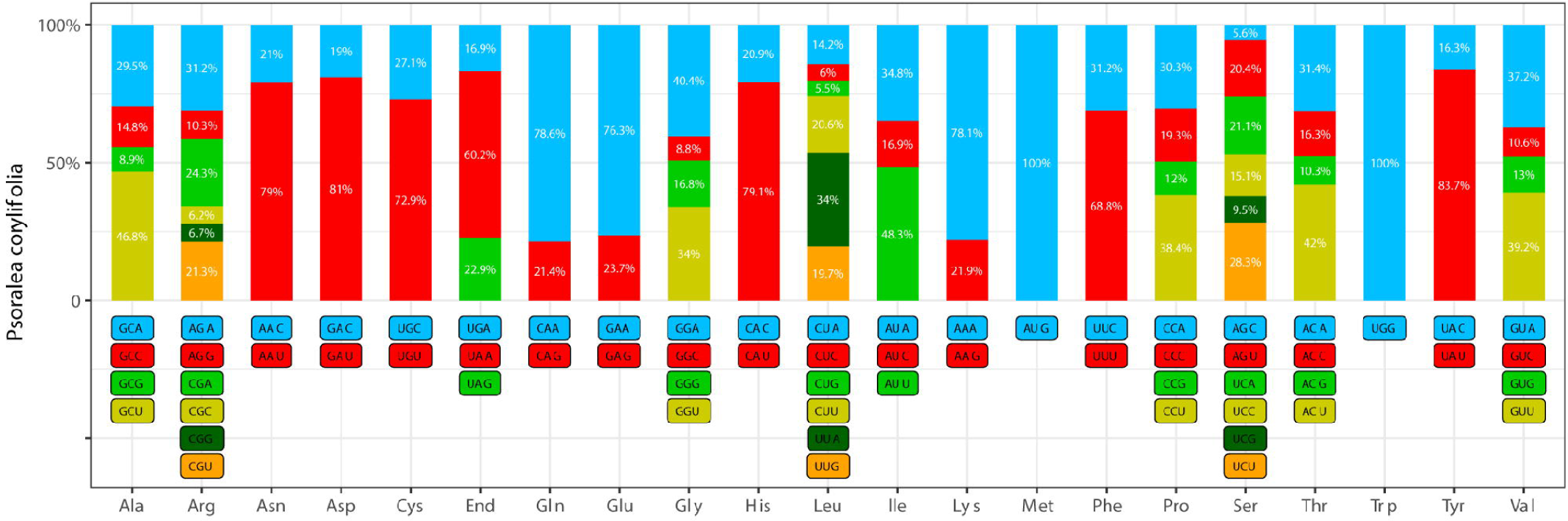
Percentage of codon usage of each amino acid in the *Psoralea corylifolia* chloroplast genome. The x-axis represents codon families.

### Repeat sequence

There existed numerous tandem repeats in the genome of eukaryotes with extremely high repetition frequency and very low order complexity. Simple sequence repeats (SSRs), consisting of 1-6 repeat nucleotide units, are frequently used as molecular markers, taking advantage of their highly reproducible, rich in polymorphism, co-dominance, and highly reliable[46]. We detected a total of 98 SSRs in *P. corylifolia* including 63 repeats for mononucleotide, 22 repeats for dinucleotide, 2 repeats for trinucleotide, 11 repeats for tetranucleotide, respectively. Because almost all chloroplast genomes had high AT content[47], and almost all SSR loci contained A or T, especially in the mononucleotide (Fig 5).In addition to the SSRs, we identified four type repeats in *P. corylifolia*, including forward, reverse, complement and palindromic repeats by REPuter. These repeats contained 14 forward repeats, 2 reverse repeats, 2 complement repeats, 32 palindromic repeats. The length of repeats varied between 30 and 257, even up to 25,557. Among these repeats, great majority repeats located in intergenic regions (S3 Table). As for tandem repeats, 49 repeats were identified in *P. corylifolia* chloroplast genome (S4 Table).

**Fig 5.**
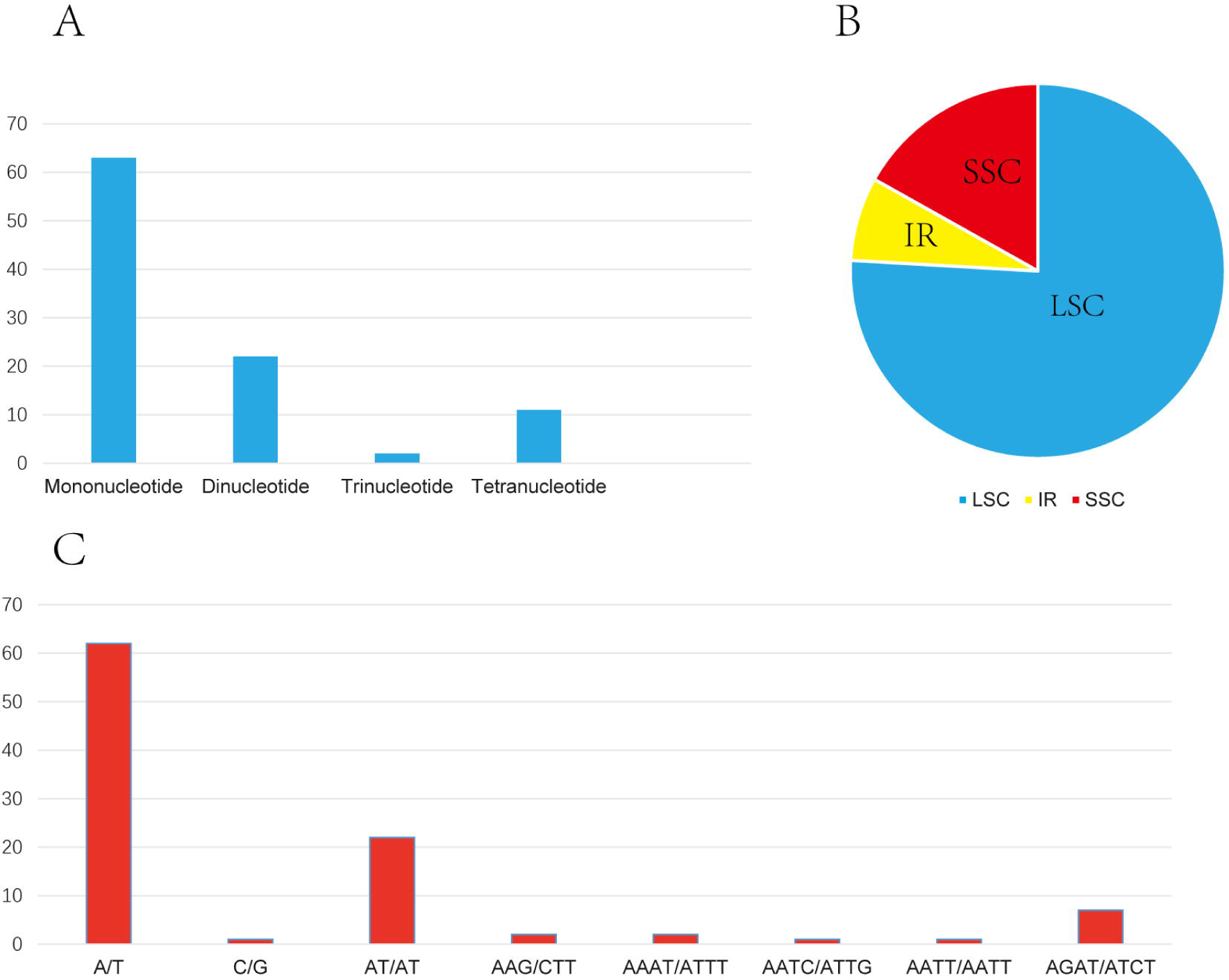
Analysis of simple sequence repeat (SSR) in the *Psoralea corylifolia* chloroplast genome. (A) Number of different SSRs types. (B) Proportion of SSRs in the LSC, IR and SSC regions, respectively. (C) Frequency of SSR loci identified in different repeat types.

### Potential RNA editing sites

It was known that RNA editing involved various procedures which could alter the protein coding sequence of a transcribed RNA by inserting, deleting or modifying nucleotides in the transcript[48]. In this study, 35 genes were employed to predict potential RNA editing sites, and 10 genes (*atpB, ycf3, atpI, atpA, psaI, petL, petG, psbB, petD, rpl2*) had been identified no sites. As a result, a total of 90 potential RNA editing sites were predicted. S to L (26%) of amino acids change occurred most, whereas P to S (2.2%) and R to C (1.1%) occurred least. Furthermore, all the type of RNA editing sites were C-to U-. We also observed that 25 (27.7%) and 65 (72.3%) editing sites were located at the first and the second codon position, respectively (S5 Table).

### Synonymous and nonsynonymous substitution rate

In genetics, dN/dS represented the ratio between a synonymous substitution (dN) and a nonsynonymous substitution (dS), which could be used to determine whether there is selective pressure on the protein-coding gene. Previous studies had estimated that, dN/dS ratios <1, =1, >1 indicating purifying selection, neutral evolution, and positive selection, respectively[49]. The *psa, psb*, and *pet* genes exhibited the lowest dN values, while the *ycf1* gene presented the highest dN values. As for dS values, the *clpP, ycf1* genes occupied the lowest and highest dS values, respectively. Furthermore, the *ccsA* and *ycf1* genes (dN/dS > 0.5) were observed to have highest the ratios of dN/dS, followed by *clpP, matK, cemA* genes (Fig 6).

**Fig 6.**
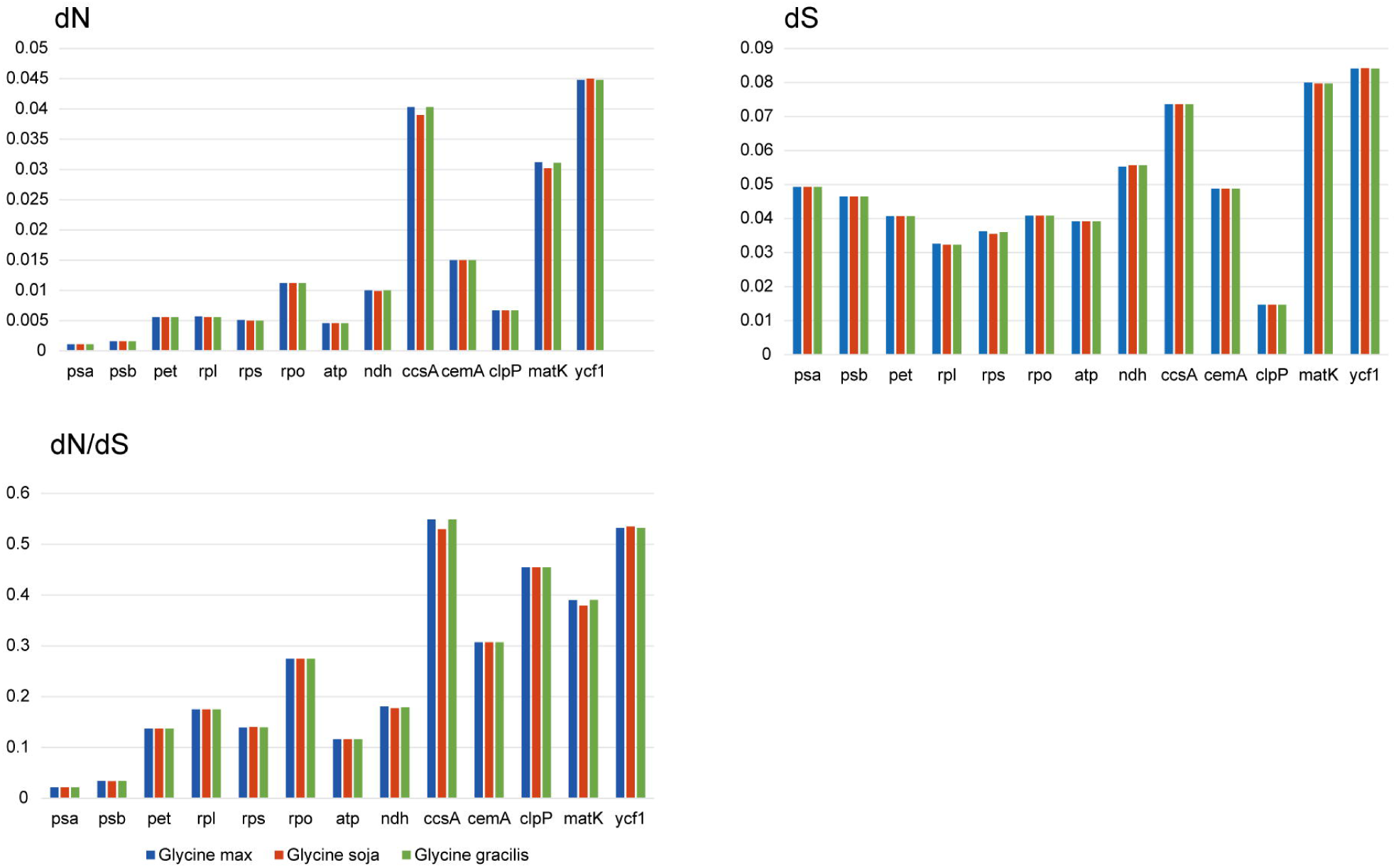
Nonsynonymous substitution (dN), synonymous substitution (dS), and dN/dS values for groups of gene or individual genes.

### Divergent Hotspots Analysis

Some highly variable regions existed in chloroplast genome sequences, which were usually clustered and described as “hotspots”[50, 51]. The nucleotide variability (Pi) was calculated by the DnaSP v6.11 software. The values ranged from 0 to 0.07028, meanwhile, the IR region was more conserved than the LSC and SSC regions. Seven mutational hotspots (Pi>0.04), including *psbD-trnT-trnE, atpI-atpH, trnG-trnS-psbI, accD-psaI-cemA, rps11-rpl36-rps8, ycf1, trnL-rpl32-ndhF*, as well as two loci (Pi>0.05) including *trnK-rbcL, rps16-accD*, exhibited remarkably higher Pi values, which was similar to the result of comparative genome analysis discovered by mVISTA program (Fig 7). What’s more, these hotspots regions would provide potential molecular marker candidates for plant identification and accelerate subsequent studies on phylogenetic analysis.

**Fig 7.**
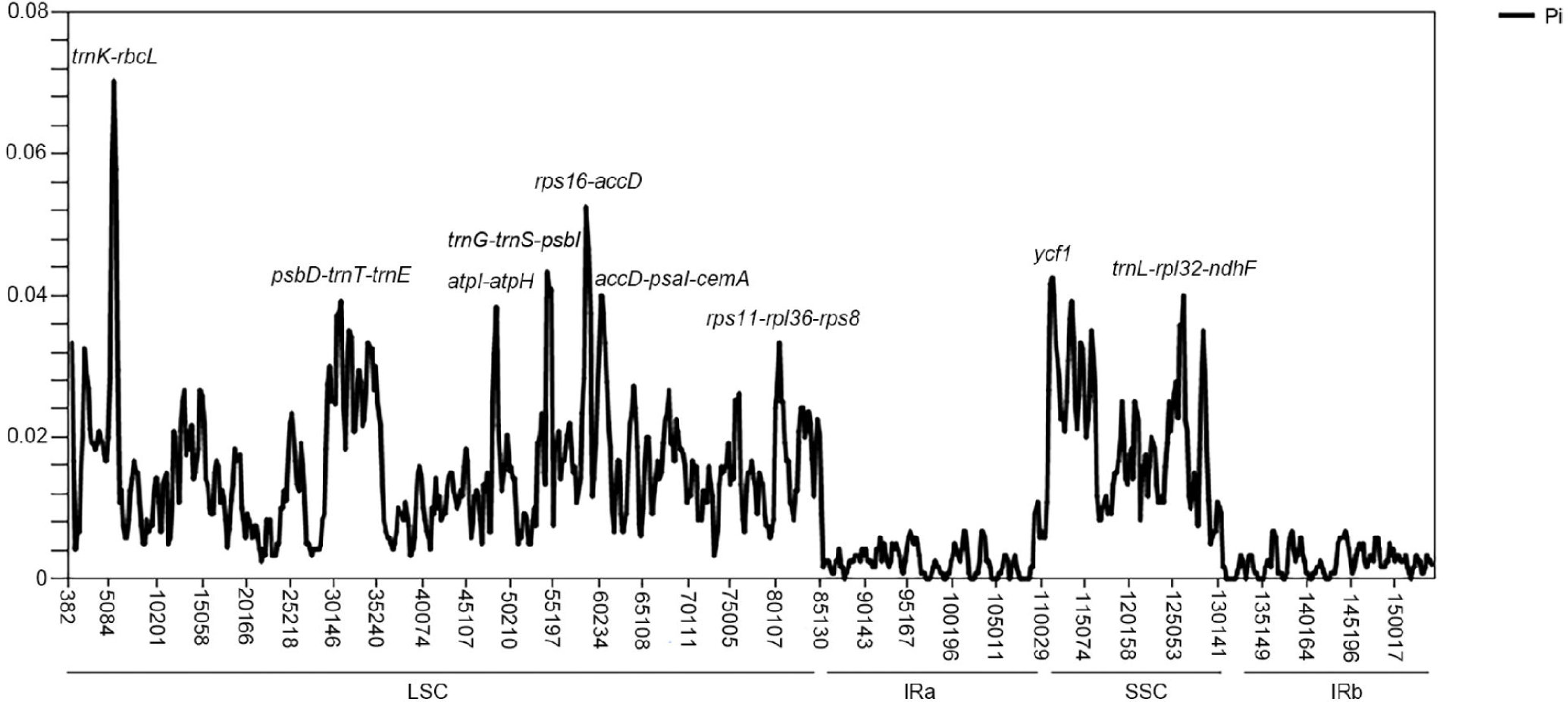
Sliding window analysis among the whole chloroplast genomes of *P. corylifolia, G. max, G. gracilis, G. soja.* The x-axis represents the position of the midpoint of a window, the y-axis represents the nucleotide diversity of each window.

### Phylogenetic and DNA barcoding analysis

We used 75 chloroplast protein-coding genes of 37 species to analyze the phylogenetic position of *P. corylifolia*. The phylogenetic trees generated by ML and BI alignment had similar topologies. In the maximum likelihood (ML) and Bayesian inference (BI) trees, almost all of nodes had a 100% bootstrap value and 1.0 Bayesian posterior probability. Furthermore, the result of *P.corylifolia* formed a single clade with three *Glycine* species strongly supported the sister relationship between *P. corylifolia* and *Glycine* species (Fig 8), which confirmed the relationship between *Psoralea* and *Glycine* genus inferred from the phylogenetic work conducted based on *trnK/matK* genes[52]. More importantly, the ML trees performed using four candidate barcodes (*ycf1, matK, accD, ndhF*) was consistent with the 75 protein-coding-based tree (S6 Fig).

**Fig 8.**
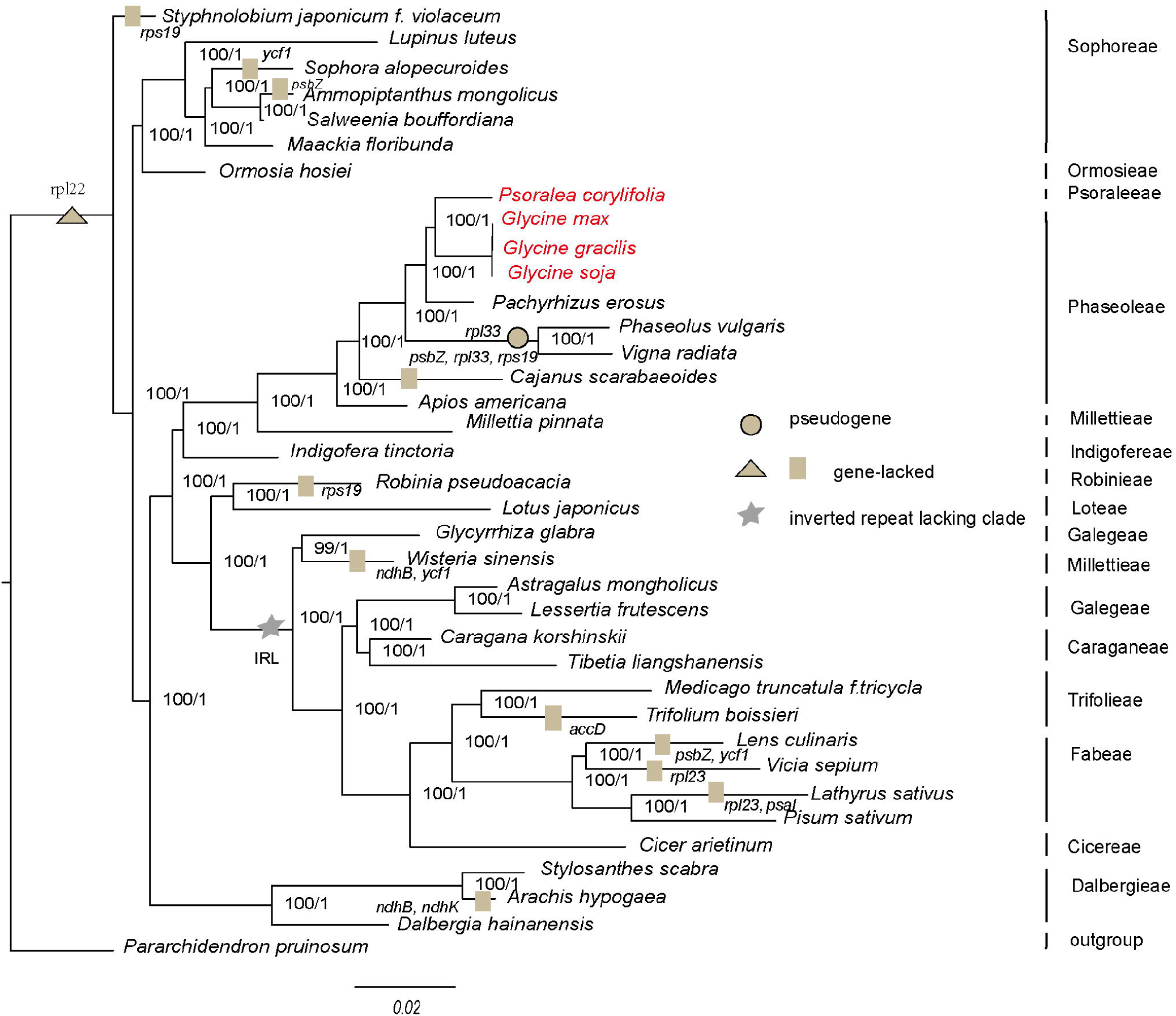
Phylogenetic trees of Papilionoideae subfamily. Phylogenetic trees reconstructed with 75 protein-coding genes of 37 species using maximum likelihood (ML) and bayesian inference (BI) methods. Numbers at nodes are values for bootstrap and Bayesian posterior probability.

### Conclusion and discussion

In this study, we present the complete chloroplast genome of *P. corylifolia* using Illumina sequencing platforms, and this is the first reported chloroplast genome assembly in the tribe Psoraleeae*. P. corylifolia* chloroplast genome (153,114 bp) is fully characterized and compare to the chloroplast genomes of related species previously reported. The chloroplast genome of *P. corylifolia* contains 111 unique genes, including 77 protein-coding genes, 4 rRNA genes and 30 tRNA genes. The ML and BI phylogenetic trees suggest that *Psoralea* is a sister genus of *Glycine*. The 16 highest divergent regions (*ycf1, matK, accD, ndhF, petD, rpl16, ndhA, trnH-psbA, trnK-rbcL, trnG-psbZ, psbD-trnT-trnE, petN-trnC, trnG-trnS, trnR-trnG-trnS, ndhI-ndhG, trnL-rpl32*) are detected.

In our study, the four highly variable protein-coding genes (*ycf1, matK, accD, ndhF*) are selected as the potential markers. Previous evidence supported that *ycf1* gene was one of the core plastid DNA barcode of land plants[53]. Meanwhile, the phylogenetic relationship of Papilionoideae based on *matK* gene had been reported[54]. Moreover, not only the *ycf1* and *matK* genes but also the rest two genes show good performance in distinguishing Phaseoleae and Psoraleeae species. These genes can be taken as promising chloroplast DNA barcodes for species identification and phylogenetic studies.

Across 36 species (genome size 176,692∼121,020) of Papilionoideae subfamily, we find that all species hold 50-kb inversion. Moreover, we also find that 13 species from 6 tribes lose their IR (IRL), including tribes Galegeae, Millettieae, Caraganeae, Trifolieae, Fabeae, as well as Cicereae. Interestingly, though *Wisteria* was very similar in appearance to *Millettia* and they classified into the same tribe (Millettieae)[52], *Wisteria* and *Millettia* are not supported as monophyletic in our result. According to this phenomenon, we speculate that it might be attributed in part to the different genome structure of *Wisteria* (IRL). In addition to these findings, a distinctive characteristic of Papilionoideae is the lack of *rpl22* genes in all species. It was quite possible that the genes loss are previously functionally transferred to the nucleus, as previously noted by researchers[55]. At the same time, it was likely that the *rpl33* gene contained stop codons within its coding region, therefore the gene presented as a pseudogene in *Vigna* and *Phaseolus*[56, 57].

In China, *P. corylifolia* is a vital traditional Chinese medicine. We believe that the complete chloroplast genome can provide helpful information for further research. The selected candidate barcodes (*ycf1, matK, accD, ndhF*) from the highest divergent regions are able to enrich the barcode library and increase the efficiency of species identification, and it is a big step forward.

## Acknowledgments

We are thankful to the National Natural Science Foundation of China (Grant no. 81673826) for its financial support to our study. We also acknowledge Ms. Liran Sun for her previous work.

## Author Contributions

**Conceptualization:** Xiaoxuan Tian, Wei Tan, Kun Zhou.

**Data Curation:** Wei Tan, Han Gao.

**Formal Analysis:** Wei Tan, Han Gao, Huanyu Zhang.

**Funding Acquisition:** Kun Zhou.

**Resources:** Wei Tan, Han Gao.

**Software:** Wei Tan, Han Gao.

**Supervision:** Xiaoxuan Tian, Kun Zhou.

**Validation:** Wei Tan, Han Gao, Xiaolei Yu.

**Writing – Original Draft Preparation:** Wei Tan.

**Writing – Review & Editing:** Xiaoxuan Tian, Weiling Jiang, Han Gao, Huanyu zhang.

## Supporting information

S1 Table. NCBI GenBank accession numbers used in this study.

S2 Table. Codon usage frequency and RSCU in the Psoralea corylifolia chloroplast genome.

S3 Table. Repeat sequences in the Psoralea corylifolia chloroplast genome.

S4 Table. Tandem repeat sequences in the Psoralea corylifolia chloroplast genome.

S5 Table. RNA editing predicted in Psoralea corylifolia chloroplast genome.

S6 Fig. Phylogenetic relationship between Phaseoleae and Psoraleeae based on *ycf1, matK, accD, ndhF* genes.

