## Supplementary figures and images for "The complete chloroplast genome of Chinese medicine (*Psoralea corylifolia Linn*): Molecular Structures, barcoding analysis, and phylogenetic Analysis"

### Supplementary Figure S6

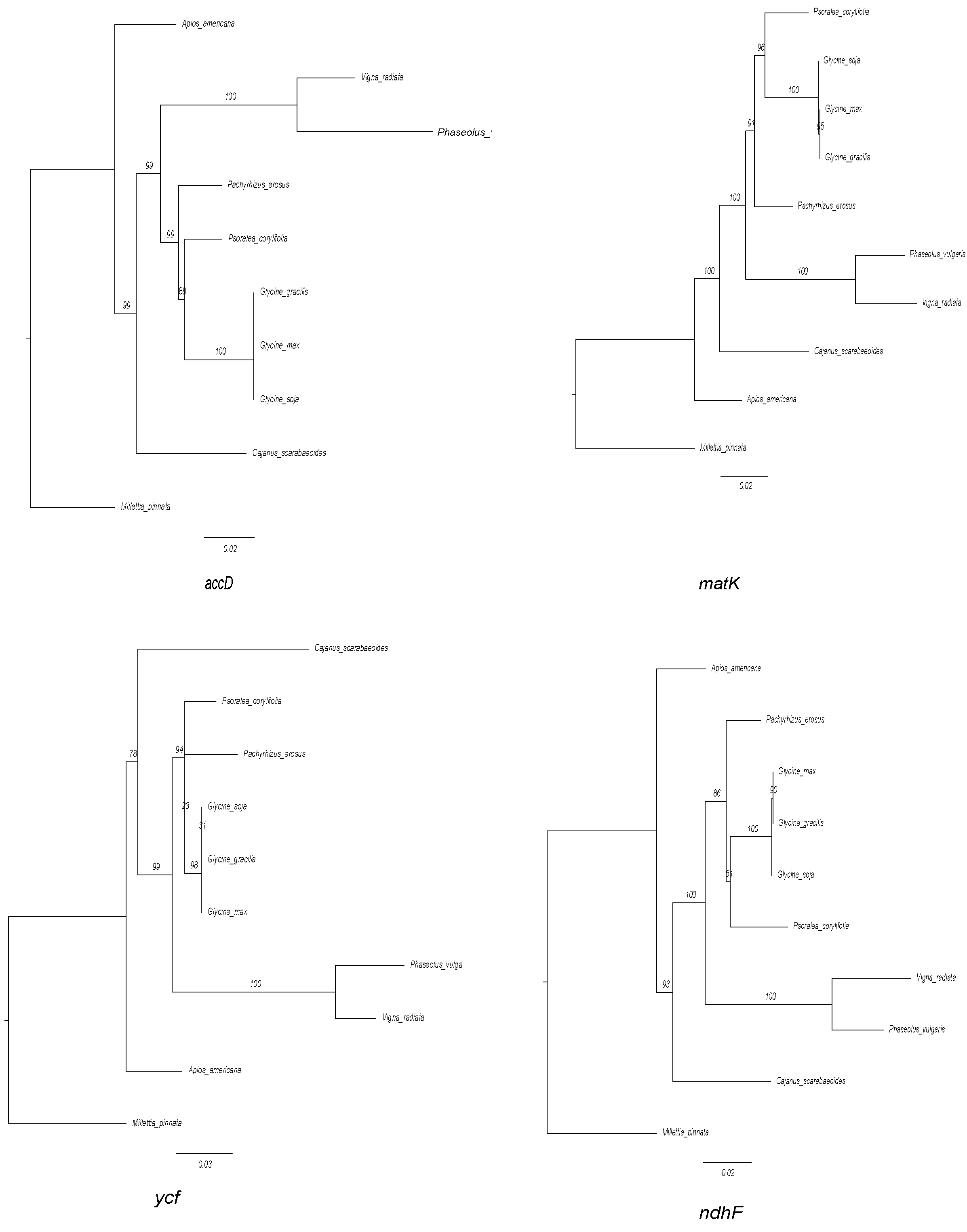
